# A selective sweep in the *Spike* gene has driven SARS-CoV-2 human adaptation

**DOI:** 10.1101/2021.02.13.431090

**Authors:** Lin Kang, Guijuan He, Amanda K. Sharp, Xiaofeng Wang, Anne M. Brown, Pawel Michalak, James Weger-Lucarelli

**Affiliations:** Edward Via College of Osteopathic Medicine, Monroe, LA, 71203, USA; School of Plant and Environmental Sciences, Virginia Tech, Blacksburg, Virginia 24061; Department of Biochemistry, Virginia Tech, Blacksburg, Virginia, USA; Research and Informatics, University Libraries, Blacksburg, Virginia, USA; Center for One Health Research, Virginia-Maryland College of Veterinary Medicine, Blacksburg, VA, 24060, USA; Institute of Evolution, Haifa University, Haifa, 3498838, Israel; Department of Biomedical Sciences and Pathobiology, Virginia Tech, VA-MD Regional College of Veterinary Medicine, Blacksburg, VA, United States of America

## Abstract

**Summary:** While SARS-CoV-2 likely has animal origins^1^, the viral genetic changes necessary to adapt this animal-derived ancestral virus to humans are largely unknown, mostly due to low levels of sequence polymorphism and the notorious difficulties in experimental manipulations of coronavirus genomes. We scanned more than 182,000 SARS-CoV-2 genomes for selective sweep signatures and found that a distinct footprint of positive selection is located around a non-synonymous change (A1114G; T372A) within the Receptor-Binding Domain of the Spike protein, which likely played a critical role in overcoming species barriers and accomplishing interspecies transmission from animals to humans. Structural analysis indicated that the substitution of threonine with an alanine in SARS-CoV-2 concomitantly removes a predicted glycosylation site at N370, resulting in more favorable binding predictions to human ACE2, the cellular receptor. Using a novel bacteria-free cloning system for manipulating RNA virus genomes, we experimentally validated that this SARS-CoV-2-unique substitution significantly increases replication in human cells relative to its putative ancestral variant. Notably, this mutation’s impact on virus replication in human cells was much greater than that of the Spike D614G mutant, which has been widely reported to have been selected for during human-to-human transmission^2,3^.

## Introduction

Severe acute respiratory syndrome coronavirus 2 (SARS-CoV-2), the causative agent of Coronavirus Disease 2019 (COVID-19), has caused over 60 million infections with at least 1.3 million deaths worldwide as of early November 2020^4^. The virus was first described in late 2019 in Wuhan, China, and quickly spread globally^1^. SARS-CoV-2 is closely related to SARS-CoV, which caused a more limited outbreak in several countries in 2003^5,6^; however, several bat and pangolin-derived viruses are even more closely related to SARS-CoV-2, indicative of a zoonotic origin ^7–9^. Bat coronavirus RaTG13—originally isolated in China from *Rhinolophus affinis* bats in 2013—shares 96% nucleotide identity with SARS-CoV-2 across the genome and ∼97% amino acid identity in the Spike (S) protein, which mediates receptor binding and membrane fusion, and is the key coronavirus determinant of host tropism^10^. Similarly, several viruses found in Malayan pangolins (*Manis javanica*) are closely related to SARS-CoV-2; with up to 97.4% amino acid concordance in the receptor-binding domain (RBD) of the S protein^8,9^. However, the exact origin and mechanism of cross-species transmission of the SARS-CoV-2 progenitor are still unknown.

In the past two decades, the emergence of severe acute respiratory syndrome coronavirus (SARS-CoV)^6,11,12^ and Middle East respiratory syndrome coronavirus (MERS-CoV)^13^ in humans and swine acute diarrhoea syndrome coronavirus (SADS-CoV) into pigs has highlighted the epidemic potential of coronaviruses^14^. Typically, only modest changes to a virus are required to initiate adaptation to a new host; for example, only two amino acid changes were necessary to produce a dramatic difference in human adaptation in both SARS-CoV and MERS-CoV S proteins^15,16^. This phenomenon is readily observed in other viruses: Ebola viruses’ human adaptation following spill-over from bats was at least partly mediated by a single alanine-to-valine mutation at position 82 in the glycoprotein^17,18^. Similarly, individual amino acid changes have been associated with recent outbreaks of several RNA viruses: chikungunya virus^19^, West Nile virus^20,21^, and Zika virus^22^. While an individual mutation that likely increases replication of SARS-CoV-2 in humans has been identified—a single aspartic acid to glycine change at position 614 in the S protein^2,3^—this occurred after emergence into humans, and the genetic determinants of SARS-CoV-2’s expansion from an animal reservoir into humans remain entirely unknown.

For a virus recently acquired through a cross-species transmission, rapid evolution, and a strong signature of positive selection are expected. For example, several rounds of adaptive changes have been demonstrated in SARS-CoV genomes during the short SARS epidemic in 2002–2003^23,24^. However, in its brief epidemic, SARS-CoV-2 has been characterized by relatively low genetic variation, concealing signals of positive selection, and leading to contradictory reports of limited positive selection^25^, “relaxed” selection^26^, or even negative (purifying) selection^27,28^. However, these results are based on dN/dS tests that are traditionally designed for eukaryotic interspecies comparisons, and thus ill-equipped to detect hallmark signatures of positive selection in viral lineages with limited sequence divergence^29^. Here, we employ highly sensitive methods enabling detection of selective sweeps, in which a selectively favorable mutation spreads all or part of the way through the population, causing a reduction in the level of sequence variability at nearby genomic sites^30^. With unprecedented statistical power that leverages information from more than 182,000 SARS-CoV-2 genomes, we demonstrate that positive selection has played a critical role in the adaptive evolution of SARS-CoV-2, manifested as selective sweeps in *Spike* and several other regions, also providing candidate mutations for further analysis and interventions. Given its role in coronavirus host tropism, we hypothesized and experimentally validated that the selective sweep identified in the S protein involves an adaptive mutation increasing replication in human lung cells, which, in turn, could facilitate more efficient human-to-human transmission.

## Results

### Selective sweeps analysis identified a Spike region with high confidence from 182,792 sequences

OmegaPlus^31^ and RAiSD^32^ were used to find putative selective sweep regions in 182,792 SARS-CoV-2 genomes downloaded from the publicly available GISAID EpiCov database (www.gisaid.org). Eight selective sweep regions were detected, including four in ORF1ab and four in the Spike region (Fig. 1 & Table 1). The Spike protein plays an important role in the receptor recognition and cell membrane fusion process during viral infection, and this protein is highly conserved among all coronaviruses. Next, we screened genomic sites in the *Spike* region that may be involved in the adaptive evolution of SARS-CoV-2 in the new host by comparing the non-synonymous differences between SARS-CoV-2 and four other *Sarbecovirus* members (one pangolin coronavirus and three bat coronaviruses; see Materials and Methods). A total of six such sites were identified (Supplementary Table 1); notably, only a single site (A1114G) was centrally located in one of the sweep regions, whereby the 372^nd^ amino acid threonine in the Spike protein of the four *Sarbecovirus* members was substituted with alanine (Thr372Ala) in human SARS-CoV-2. Out of the 182,792 SARS-CoV-2 genomes, no sequence polymorphism was found in this position (1114G), suggesting a rapid fixation of this mutation via hard sweep. The alternative, putatively ancestral, coronavirus variant (A1114) was perfectly conserved in *Sarbecovirus* members from bats and pangolin.

**Fig 1.**
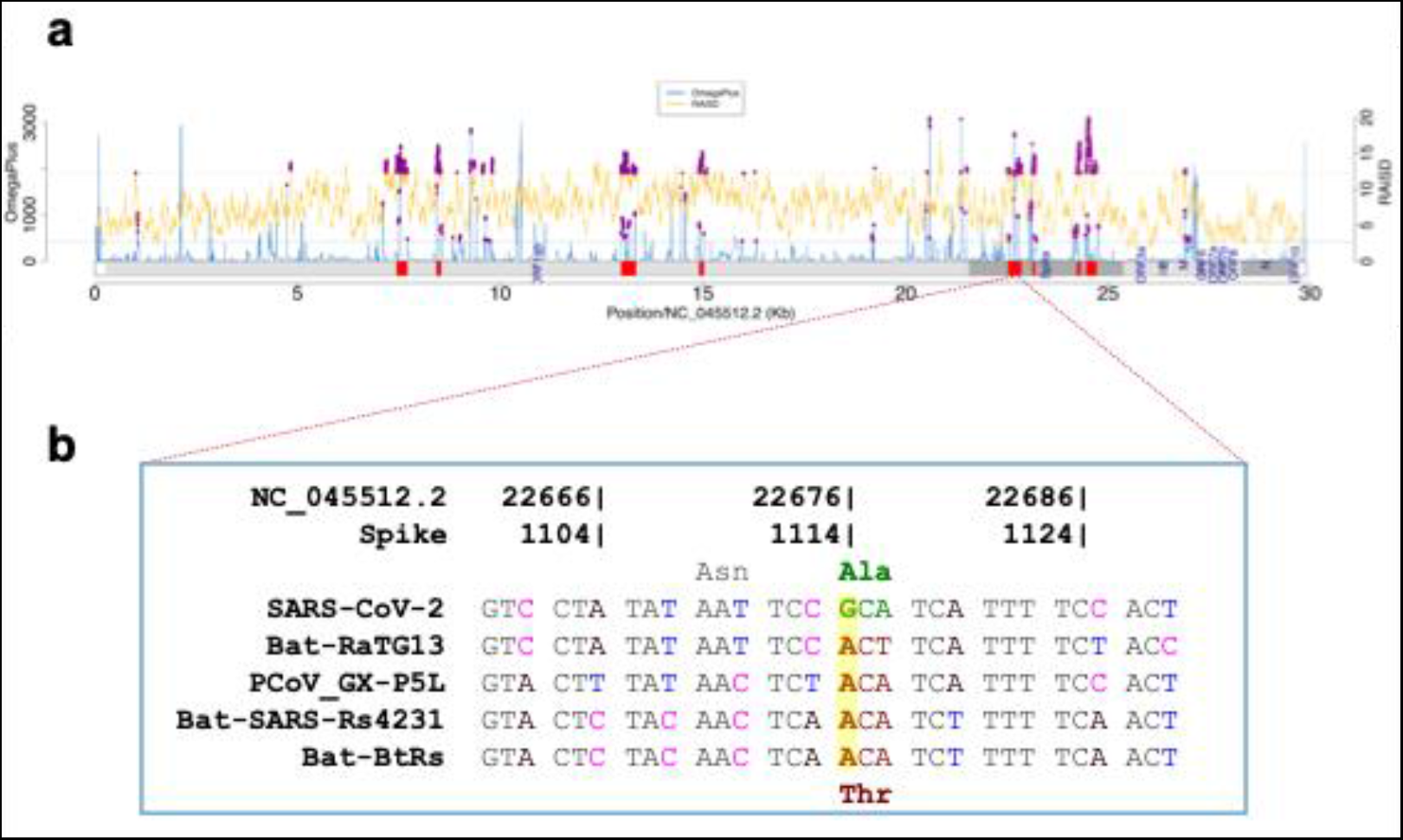
Selective sweeps analysis. (a) Selective sweep regions (shown as red blocks) identified in 182,792 SARS-CoV-2 genomes, using OmegaPlus (blue lines) and RAiSD (yellow lines). The common outliers (0.05 cutoff, purple dots) from the two methods were used to define selective sweep regions. (b) Non-synonymous difference (Thr372Ala) between SARS-CoV-2 and four other *Sarbecovirus* members found in the putative selective sweep region (22,529-22,862).

**Table 1:** Putative sweep regions (the region containing the Spike G1114A position is bolded).

| Start | End | Region |
| --- | --- | --- |
| 7,445 | 7,711 | ORF1ab |
| 8,426 | 8,542 | ORF1ab |
| 12,978 | 13,350 | ORF1ab |
| 14,907 | 15,027 | ORF1ab |
| **22,529** | **22,862** | **Spike** |
| 23,132 | 23,196 | Spike |
| 24,225 | 24,319 | Spike |
| 24,456 | 24,712 | Spike |

### Structure-based analysis of SARS-CoV-2 S protein variants

Comparative molecular modeling of WT (372A - SARS-CoV-2), 372T, and 614G S protein was performed to connect the selective sweep G1114A mutation to structural data (Fig. 2). Structures were energy-minimized after mutation and analyzed for change in ACE2 binding and probability of N-linked glycosylation sites as a result of mutation. Increased probability of N-linked glycosylation at N370 was observed in the 372T variant (Fig. 2a-c), given the mutation to a threonine both provided a standard N-linked glycosylation site motif (NXT/S) and the solvent accessible surface area of N370 (Fig. 2d-e). No glycosylation site was predicted at N370 in the WT S protein. To further probe the impact of the predicted glycosylation site at N370 as a result of presence of a threonine at position 372 of S protein, N370 was glycosylated with a mannose glycan and energy-minimized on the 372T S protein model in order to observe any minor side chain readjustment as a result of N-glycan presence. N370 glycosylation of 372T S protein occurs in close structural proximity to the essential glycosylation site of N343^33^, further providing additional N-glycans shielding of the RBD (Fig. 2a). Surface maps also reveal an additional space-filling and polar surface that is now occupied by the N370 N-glycan (Fig. 2b-c). Additionally, Molecular Mechanics Generalized Born Surface Area (MM/GBSA) free energy of binding of ACE2 to S protein was calculated for WT, 372T, and N370-glycosylated 372T S protein. Free energy of binding of ACE2 to WT S protein showed a very negative, favorable relative binding affinity (−180.503 kcal/mol) while free energy of binding of ACE2 to the putatively ancestral 372T variant and N370-glycosylated 372T variant was less negative and favorable (−95.7685 kcal/mol and -69.825 kcal/mol, respectively), highlighting that while the glycosylation at N370 is not in close proximity (>10 Å) of the receptor binding motif (RBM), it is influencing the RBD of S protein and its potentiality in binding ACE2. Structural analysis of 614G did not indicate any major modifications to the structure of S protein. Residue 614 is in close proximity to a glycosylation site (N616) but 614G did not change probability of glycosylation or general surface properties compared to 614D.

**Fig 2.**
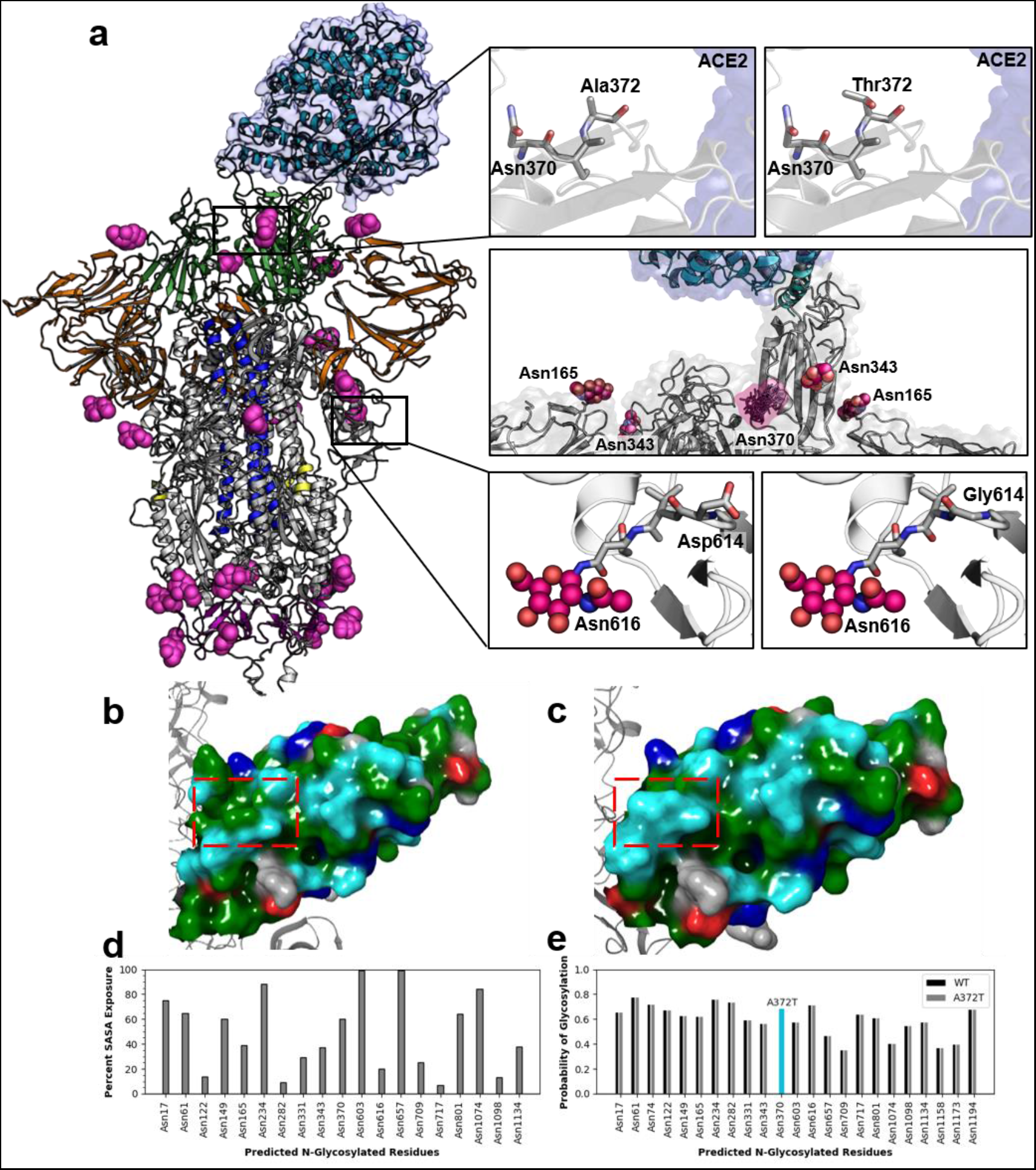
Structure-based analysis of SARS-CoV-2 S protein variants. **(a)** Visualization of 372T and D614G mutants. Structure of S Protein (PDB 7A94) is displayed in cartoon and colored by RBD (green), NTD (orange), CH (blue), FP (yellow), and CD (pink). Glycans are displayed in spheres colored in hot pink. Top panel shows the WT (372A) and 372T mutant, the middle panel displays a glycosylated N370 372T S protein with various rotamers of the mannose glycosylated N370, and the bottom panel shows the WT and 614G mutant. **(b)** Surface map of the WT S protein and **(c)** The N370-glycosylated 372T S protein, colored by the residue side-chain properties – colors represent: green for hydrophobic, blue for positively charged, red for negatively charged, teal for polar uncharged, and gray for neutral. **(d)** Predicted N-glycosylated residues identified by Schrödinger-Maestro’s BioLuminate (v. 2020-2) Reactive Residue package with percent solvent accessible surface area (SASA) exposure of each residue. **(e)** Predicted N-glycosylated residues identified by NetNGlyc 1.0 Server with the probability of being glycosylated.

### SARS-CoV-2 Spike A372T decreases replication in human cells

We generated the Spike A372T reverse mutant with the putatively ancestral G1114 nucleotide, using a bacteria-free cloning approach we have previously developed to prevent bacterial toxicity associated with manipulating unstable viral genomes in bacteria^34,35^. Concurrently, we generated the Spike D614G mutant that has been associated with higher titers in nasopharyngeal swabs in humans and increased replication in human cells and hamsters^2,3^. Both mutants were constructed in an infectious clone originally produced in yeast^36^ of SARS-CoV-2 strain 2019-nCoV BetaCoV/Wuhan/WIV04/2019^1^. A schematic of the A372T mutant is presented in Figure 3a; while not depicted, the D614G mutant was made by replacing the WT codon (GAT) with the glycine-encoding codon (GGC). Following virus rescue, viral plaque morphology was similar for all three viruses, although the A372T mutant plaques appear slightly smaller (Fig. 3b).

**Figure 3.**
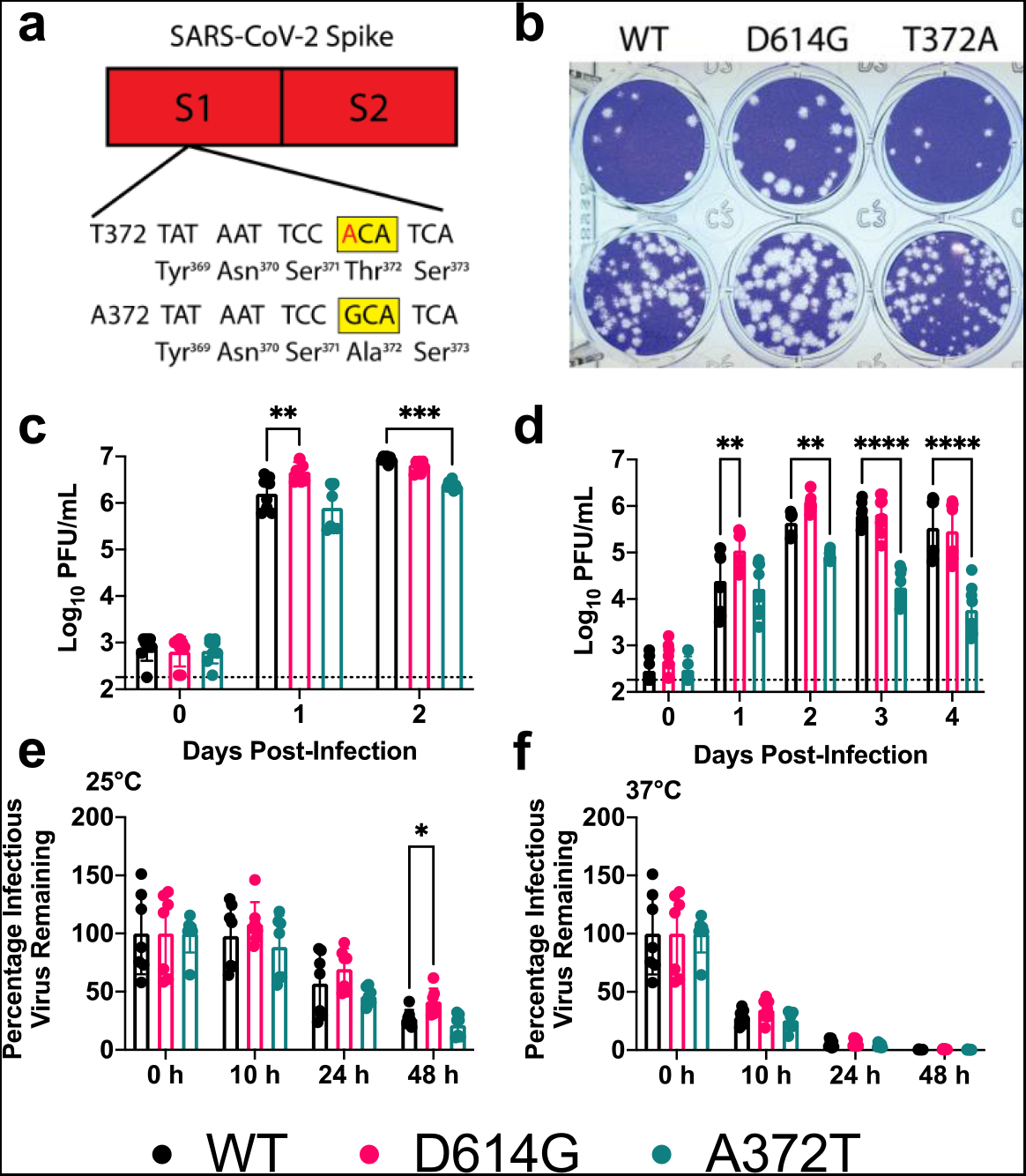
A372T Spike substitution decreases SARS-CoV-2 replication on human lung epithelial cells. **(a)** The Spike T372 SARS-CoV-2 mutant was generated by making a single G-to-A substitution. The mutant nucleotide is presented in red and the altered codon is highlighted in a yellow box. **(b)** Plaque morphology of WT and mutant viruses. Plaques were visualized 2 days post-infection (dpi) on Vero E6 cells. **(c-d)** Viral replication on Vero E6 (c) and Calu-3 cells (d) following an infection at an MOI of 0.05. The sample at 0 dpi was collected immediately after infection to ensure cells were exposed to similar levels of virus, and then samples were collected at 24 hour intervals. **(e-f)** Kinetics of thermal stability. A solution of 105 PFU of each virus was incubated at the indicated temperature for different lengths of time. Infectious virus was measured by plaque assay on Vero E6 cells. Statistical comparisons were made using two-way ANOVA with Dunnett’s multiple comparisons test; *p<0.05,**p<0.01,***p<0.001, ****p<0.0001.

We next evaluated the replication kinetics of each virus--WT, Spike A372T, and Spike D614G--in Vero E6 and Calu-3 cell lines, monkey kidney and human lung epithelial cell lines, respectively. Following infection in Vero E6 cells, viral titers rose rapidly for all three viruses, and only minor differences in peak titers were observed among the viruses (Fig. 3c). In Calu-3 cells, the D614G mutant produced significantly higher titers than WT 1 day post-infection (dpi) but levels were similar for the remaining timepoints (Fig. 3d; p=0.0066 by 2way ANOVA with Dunnett’s correction at 1 dpi). No differences were observed 24 hours after infection between the WT and A372T mutant but later timepoints showed a marked reduction in replication for the A372T mutant (p=0.0033, <0.0001, and <0.0001 for 2, 3, and 4 dpi, respectively). Compared to WT, the D614G had modest differences of 2.9-, 2.9-, 1.3-, and 0.8-fold in viral titers on 1, 2, 3, and 4 dpi, respectively; in contrast, as compared to WT, A372T titers were 1.8-, 5.5-, 31.1-, and 64.1-fold lower on 1, 2, 3, and 4 dpi, respectively (Fig. 3d). These data indicate that an alanine at Spike position 372 confers a robust fitness advantage over several timepoints in human lung cells and that this effect is considerably more substantial than the change at position 614.

Based on structural analysis, others postulated that the SARS-CoV-2 Spike trimer would have higher thermal stability than the Spike of bat virus RaTG13^37^. To determine if A372T altered SARS-CoV-2 thermal stability, we incubated 10^5^ PFU of WT SARS-CoV-2, D614G, or A372T at room temperature (∼25°C) or 37°C to mimic environmental and human body temperature, respectively. A372T titers did not significantly differ from WT at any timepoint for either temperature (Fig. 3e-f). Following 48-hours incubation at room temperature, the titer of D614G was higher than WT SARS-CoV-2 (p=0.0303), which is consistent with previous reports^3^.

## Discussion

COVID-19 has now claimed the lives of >2 million people worldwide, dwarfing the number of deaths caused by SARS-CoV (774^38^) or MERS-CoV (858^39^). Although phylogenetic and epidemiological data suggest a zoonotic origin for SARS-CoV-2, little is known about the viral mutations that likely occurred to adapt the virus to human transmission. The SARS-CoV-2 progenitor would have likely required new adaptations to sustain human-to-human transmission—a process that likely included a strong positive selection event, favoring the viruses with the greatest replication in the human respiratory tract. Here, we identified a region in the *Spike* gene with a strong signal of such an event—a selective sweep—from over 180,000 SARS-CoV-2 genomes. Within this region, present in the receptor-binding domain, we identified a non-synonymous single nucleotide polymorphism (SNP) that is fixed in all SARS-CoV-2 genomes sequenced to date, while an alternative, and presumably ancestral SNP, is fixed in the other members of the *Sarbecovirus* lineage.

Residue 372 lies within the RBD (Fig. 2a), which mediates viral entry through the human ACE2 receptor^1^. While positioned adjacent to the ACE2 interface of the RBD, the presence of an alanine at position 372 (372A) is predicted to remove a glycosylation site present at the asparagine at position 370^37^, which may alter S protein maturation or receptor binding (Fig. 2). Indeed, molecular modeling of an N-glycan at N370 in an open conformation of 372T S protein shows a highly solvent accessible glycan site (Fig. 2). In the closed conformation of 372T, the N370 glycan site becomes less solvent exposed and further fills a solvent-accessible region on the outer edge of the RBD. N-Glycans are known to modulate the RBD of S protein, with glycans at position N165 and N234 influencing the open/closed metastable conformation states of the RBD, and N-glycans at N331 and N343 serving more of a shielding role of the RBD itself regardless of state^33^. N370 glycosylation is in close structural proximity of the N-glycan site at N343 and is in relative distance to the RBM and RBD/ACE2 interface (Fig. 2a-c). Free energy of binding of ACE2 to S protein indicates a decrease in relative binding affinity of ACE2 to S protein in the N370-glycosylated 372T variant compared to WT (−69.825 kcal/mol vs -180.503 kcal/mol, respectively). We hypothesize that while an additional N-glycan at N370 in 372T S protein could contribute to glycan shielding of the RBD, its proximity to the RBM and center positioning between essential glycan sites could preclude RBD/ACE2 interaction affinity or ability to transition between open/close states, decreasing RBD/ACE2 affinity and subsequent virulence.

Using a reverse genetics system to generate a SARS-CoV-2 mutant containing the putative ancestral SNP, we show that the A372T S mutant virus replicates over 60-fold less efficiently than WT SARS-CoV-2 in Calu-3 human lung epithelial cells (Fig. 3d). Further, the growth of the A372T S mutant was greatly reduced for multiple days, which may be indicative of an impact on viral shedding kinetics in humans. Of note, we also generated the D614G S mutant here--widely reported to increase SARS-CoV-2 infectivity^2^—which only increased viral titers by a maximum of 2.9-fold in Calu-3 cells compared to WT, a finding that is consistent with previous results^3^. We also observed a slight attenuation for the A372T S mutant in Vero E6 cells (3.8-fold lower titers compared to WT 2 dpi). The large differences in replication differences between the two cell lines suggest a cell-specific mechanism of attenuation. In fact, besides their species of origin, Calu-3 and Vero E6 cells differ in several important aspects. First, Vero E6 cells are deficient in type-1 interferon signaling^40^, which inhibits SARS-CoV-2 replication^41,42^. However, the S protein is not known to antagonize IFN production, and, therefore, interferon is unlikely driving the differences observed here. Additionally, the S protein requires host-mediated proteolytic cleavage to undergo fusion, which can be driven by several proteases, including TMPRSS2 at the cell surface and cathepsins B and L (CatB/L) in endosomes^43^. Notably, Calu-3 cells express low levels of cathepsins but high levels of TMPRSS2, suggesting a TMPRSS2-dependent entry mechanism in Calu-3 cells^44^. In contrast, SARS-CoV-2 infection of Vero E6 cells is CatB/L dependent^43^. Clinical isolates of coronaviruses prefer entry through TMPRSS2 as opposed to CatB/L^45,46^; accordingly, Calu-3 cells mimic the human environment closely in terms of S protein priming. Thus, these data hint that the A372T S mutant’s attenuation could be mediated by inefficient TMPRSS2 cleavage of Spike. Host proteases have been implicated in the cross-species transmission of MERS-CoV from bats to humans^16,47^. Hence, it will be important for future studies to define the importance of TMPRSS2-mediated cleavage of the S protein in the context of these mutations.

We did not observe large temperature stability differences between viruses here. A previous report predicted that the SARS-CoV-2 S protein would have higher thermal stability than the S protein of bat coronavirus RaTG13^37^; however, it does not appear that the residue difference at position 372 dictates this difference. It may also be that at different timepoints or temperatures that differences would have been observed; nonetheless, these data suggest that thermal stability is not a likely driving factor in the emergence of the variants at position 372 or 614 in the S protein.

Overall, our data supply solid evidence that S protein residue 372 is critical for replication in human cells. The fact that this site is not polymorphic in >180,000 SARS-CoV-2 sequences further underscores its importance. The threonine-to-alanine change may have enabled the putative ancestral virus to replicate more efficiently in human cells, thus enabling efficient human-to-human transmission. While other studies have identified evidence of positive selection in SARS-CoV-2^2,25,48^, these studies are either entirely computational or use pseudotyped viruses. Although useful information can be obtained using pseudotyped viruses, they typically express only the Spike protein; consequently, they do not fully recapitulate the viral life cycle, including interactions between different viral proteins and the host, and cannot complete an entire viral replication cycle. Plante et al. used a reverse genetics system to generate the D614G S protein mutant and showed increased replication in cell culture and hamsters ^3^, highlighting the utility of using live virus to characterize critical viral mutations. Our use of live virus enables future studies in hamster or ferret models that recapitulate human-to-human transmission^49–52^.

Although the experimental data presented here clearly demonstrate the dramatic impact of the S protein A372T mutation on virus replication in human lung cells, we cannot definitively conclude that it enabled efficient human-to-human transmission or that it was necessary for cross-species transmission. Our findings suggest, though, that efficient replication in a human would be unlikely with a threonine at S protein position 372, from which we could infer that transmission would be equally unlikely. Since the true putative SARS-CoV-2 ancestor has not been isolated, it is impossible to know when this mutation may have arisen. Phylogenetic estimates suggest SARS-CoV-2 emerged in late November 2019 to early December 2019^53^, though the first known case was not detected until December 1st, 2019^54^. However, this case had no connection to the Huanan seafood market, indicating that transmission was ongoing before early December or that the seafood market is not the origin of the pandemic, but rather a spreading point. While it is impossible to know SARS-CoV-2’s exact emergence date, it seems likely that transmission occurred unnoticed for some period of time, thereby providing a window for SARS-CoV-2’s ancestor to adapt to human replication.

## Materials and Methods

### Ethics and biosafety

The generation of recombinant SARS-CoV-2 was approved by the Institutional Biosafety Committee at Virginia Tech. All studies with live infectious SARS-CoV-2 or mutant viruses were performed in an approved BSL3 facility following CDC and NIH guidelines. Researchers manipulating live virus wore an N95 respirator or Powered Air Purifying Respirators (PAPR) as approved by the IBC.

### Putative Selective Sweep Region Detection

A total of 182,792 complete SARS-CoV-2 genomes (low coverage genomes with N’s > 5% were excluded) were downloaded from the GISAID EpiCov database (www.gisaid.org) as of Nov. 11, 2020. Sequences were first aligned to SARS-CoV-2 reference (NCBI Reference Sequence/NC_045512.2) using Minimap2^55^ (with default parameters other than ‘-ax asm5’). Sequences with aligned lengths less than 20,000 were excluded from the analysis. The 136,114 remaining sequences were then aligned by using MAFFT^56^. OmegaPlus^31^ and RAiSD^32^ were used for sweep region detection, and the bat coronavirus RaTG13 (GenBank/MN996532.2) genome was used as an outgroup. OmegaPlus was performed with parameters ‘-minwin 100 -maxwin 1000 -grid 20000 -impute’. RAiSD was executed with parameters ‘-c 1 - w 50 -M 1 -y 1 -G 20000 -COT 0.05 -O -COD 100’. The common-outlier method integrated in RAiSD was used to identify the overlapped positions reported by both methods, with setting the cut-off threshold of 0.05 (−COT 0.05) and the maximum distance between outliers of 100 (−COD 100). Finally, the common outliers were manually grouped into eight regions with the size of each region greater than 50 bp. Four genome sequences (Pangolin coronavirus isolate PCoV_GX-P5L: GenBank/MT040335.1; Bat coronavirus RaTG13: GenBank/ MN996532.2; Bat SARS-like coronavirus isolate Rs4231: GenBank/KY417146.1; Bat coronavirus BtRs-BetaCoV: GenBank/MK211376.1) were used to assess the nucleotide changes among different Sarbecovirus members.

### Molecular Modeling and Free Energy of Binding Calculations

Glycosylated S Protein structure was downloaded from the RCSB Protein Data Bank (PDB ID: 7A94^57^) and was energy minimized using Schrödinger-Maestro (v. 2020-2) software^58^. The S Protein was mutated using PyMOL^59^ to the D614G and A372T S protein variants. After mutation, energy minimization was performed using the OPLS3e force field. To identify glycosylation propensity and predicted glycosylated residues of the WT S Protein and the A372T mutant, the NetNGlyc 1.0 Server^60^ and Schrödinger-Maestro’s BioLuminate (v. 2020-2) Reactive Residue package was used^58,61–63^. Schrödinger-Maestro’s (v. 2020-2) Workspace Operations was used for glycosylation of the Asn370 with mannose, to identify various Asn370-glycan rotamers, to analyzes the surface residue properties of the WT S Protein and A372T mutant. Molecular Mechanics/Generalized Born Surface Area (MM/GBSA) binding free energy was calculated using Schrödinger-Maestro’s (v. 2020-1) Prime Package^64,65^. Structures were visualized using PyMOL.

### Cell lines

Vero E6, monkey kidney cells (ATCC CRL1586) and Calu-3, human lung epithelial cells (ATCC HTB-55) were purchased from ATCC. Vero E6 were maintained in Dulbecco’s Modified Eagle’s medium (DMEM) containing 5% fetal bovine serum (FBS; R&D Systems), gentamicin (50 μg/mL), 10 mM HEPES, and 1x nonessential amino acids (NEAA). Calu-3 were maintained in DMEM with the same additives except with 20% FBS. All cell lines were maintained in a humidified incubator at 37°C.

### Virus strains

Infectious SARS-CoV-2 strain 2019-nCoV BetaCoV/Wuhan/WIV04/2019 was recovered from an infectious clone originally described by Thao et al.^36^. The viral rescue procedure is described below.

### Nucleoprotein expression construct

Homologous recombination was used to make the N gene expression plasmid in yeast. The Saccharomyces cerevisiae strain YPH500 (MATα ura3-52, lys2-801, ade2-101, trp1-Δ63, his3-Δ200, leu2-Δ1) was used for homologous recombination. All yeast cells were grown at 30°C in a synthetic defined (SD) medium containing 2% glucose as the carbon source. Histidine was omitted from the growth medium to maintain plasmid selection. To construct the nucleoprotein expression vector, pRS313 was linearized by digestion with BamHI and XbaI to serve as the backbone. The PCR-amplified N gene product was cloned into pRS313 by homologous recombination in yeast cells. Plasmid DNA was extracted from the yeast colonies that grew on SD-His plate and then transformed into *E. coli* for amplification. The construct pRS313-T7-N was sequencing-confirmed with the correct coding sequence of the N gene and expected junction sites where the N gene is inserted downstream of the T7 promoter and upstream of the EcoRI site. The primers used for cloning were: Forward: 5’ gtaaaacgacggccagtgaattgtaatacgactcactatagATGTCTGATAATGGACCCC 3’ and Reverse: 5’ cctcgaggtcgacggtatcgataagcttgatatcgaattcTTAGGCCTGAGTTGAGTCAG 3’ where uppercase letters represent the sequences of N gene and lowercase letters are from vector sequences.

### Bacteria-free cloning (BFC) and site-directed mutagenesis (SDM)

SDM was performed using BFC, starting with the yeast clone as a template. Primers for mutagenesis were obtained from Integrated DNA Technologies (IDT). PCRs were performed using Platinum SuperFi PCR Master Mix (Invitrogen) or repliQa HiFi ToughMix (Quantabio). Amplicons were purified from a GelGreen nucleic acid stained-gel (Biotium) using the NucleoSpin Gel and PCR clean-up kit (Macherey-Nagel). Gel-purified amplicons were then assembled using NEBuilder HiFi DNA Assembly Master Mix at a 1:1 molar ratio for each DNA fragment and incubated at 50°C for two hours. To confirm that no parental yeast-clone was carried through the process, we included a control containing the DNA fragments but no assembly mix; this was then treated identically to the other samples for the remainder of the process. The assembly was then digested with Exonuclease I, Exonuclease III, and DpnI to remove single-stranded DNA, double-stranded DNA, and bacterial-derived plasmid DNA, respectively; note, in this case, DpnI was not strictly necessary because yeast-derived plasmids are resistant to DpnI-cleavage^66^; however, it was included for consistency with our previous studies^34^. We then amplified the circular product using the FemtoPhi DNA Amplification (RCA) Kit with Random Primers (Evomic Science).

### Virus rescue

RCA reactions were linearized with EagI-HF (NEB) and then column purified (Macherey-Nagel). The N expression plasmid was linearized using EcoRV-HF (NEB). Capped-RNA was produced using the mMESSAGE mMACHINE T7 Transcription Kit (Invitrogen) by overnight incubation (∼16 h) at 20°C using 2-3 μg of DNA. We used this lower temperature to obtain more full-length transcripts ^67^. Reactions for full-length viral transcripts were supplemented with an additional 4.5 mM of GTP.

We electroporated the RNA transcripts into a mixture of Vero E6 (75%) and BHK-21 (25%) cells containing a total of 2×10^7^ cells per electroporation^36^. The Bio-Rad Gene Pulser Xcell Electroporation System was used with the following conditions: 270 volts, resistance set to infinity, and capacitance of 950 μF^68^. Before pulsing, the cells were washed thoroughly and then resuspended in Opti-Mem (Invitrogen). Following a single pulse, cells were allowed to incubate at room temperature for 5 minutes and we then added fresh growth media before seeding a T-75 flask and placing it at 37°C with 5% CO_2_. The cells were monitored daily and the supernatant was harvested at 25% CPE. Sequences were confirmed by Sanger sequencing of virus stocks. Only virus direct from transfection (p0 stock) was used for further characterization. Virus titers were assessed by plaque assay on Vero E6 cells.

### Plaque assays and growth curves

Viral titration was performed on Vero E6 cells by plaque assay. Briefly, serial ten-fold dilutions of each sample were made and then added to confluent monolayers of Vero E6 cells. An overlay containing 0.6% tragacanth gum (Millipore Cat# 104792) was then added; plaques were visualized following formalin fixation and staining with crystal violet. For growth curves, Vero E6 and Calu-3 were infected at a multiplicity of infection (MOI) of 0.05 with each virus. Following one-hour of infection, we removed the virus inoculum, washed once with 1x PBS, and added fresh growth media. We then collected supernatant as the 0-day timepoint and daily thereafter until 50% cytopathic effect (CPE) was observed, each time replacing the volume taken with fresh growth media. Infectious virus was measured by plaque assay on Vero E6 cells.

### Temperature stability

A virus stock containing 10^5^ PFU of each virus was prepared in RPMI-1640 containing 2% FBS and 10 mM HEPES. The virus stock was aliquoted into tubes in triplicate or quadruplicate for each timepoint; a 0-hour timepoint was collected immediately and stored at -80°C for normalization. At each timepoint, we placed a subset of the tubes at -80°C for storage until virus titration by plaque assay. The remaining virus was calculated by dividing the individual titers at each timepoint by the average of the viral titer at the 0-hour timepoint.

### Statistics

Statistical analyses were performed in GraphPad version 9. Viral titers were compared to WT using a two-way ANOVA with Dunnett’s multiple comparisons test; a *p-value* of less than 0.05 was considered significant. The limit of detection for our plaque assays is 2.3 log_10_ PFU/mL; however, negative values were given an arbitrary value of 0.9 plaques for a ten-fold diluted sample, which corresponds to 2.26 log10 PFU/mL.

## Acknowledgements

This work was supported by a Center for One Health Research Seed Program grant awarded to P.M. and J.W-L. and an Institute for Critical Technology and Applied Science Junior Faculty Award to J.W-L. We thank Dr. Volker Thiel for sharing the yeast SARS-CoV-2 reverse genetics platform.

## Author contributions

Conceptualization: L.K., P.M., J.W-L.; Investigation: L.K., G.H., A.K.S., A.M.B., P.M., J.W-L.; Writing-Original Draft, L.K., A.M.B., P.M., J.W-L.; Writing-Review &Editing, L.K., X.W., A.M.B., P.M., J.W-L.; Supervision, X.W., A.M.B., P.M., J.W-L.; Funding Acquisition: P.M., J.W-L.

